# A Symphony of Genres: Driving Information Dynamics in Functional Brain Networks

**DOI:** 10.64898/2026.04.22.720162

**Authors:** Abolfazl HaqiqiFar, Azin Shirmohammadi, Amirhossein Yekta, G. Reza Jafari

**Affiliations:** Department of Physics, Shahid Beheshti University, Evin, Tehran, Iran; Center for Communications Technology, London Metropolitan University, London N7 8DB, UK

**Keywords:** EEG, Transfer Entropy, Rich-Club, Directed Connectivity, Music Cognition

## Abstract

While the influence of music on brain networks is well-established as a behavioral and cognitive phenomenon, how different musical genres modulate these networks remains an open question. In this study, we analyzed a publicly available 128-channel EEG dataset recorded from 20 participants during eyes-closed listening to 12 music genres. To investigate genre-specific effects, we constructed directed information flow networks using transfer entropy. The backbone of these networks was then extracted using a two-stage pipeline comprising sparse encoder auto-denoising followed by an entropy-based inter-subject stability filter. Our analysis revealed that music listening significantly alters information routing within brain networks. Critically, different musical genres induced distinct patterns of neural integration. Rhythmically and structurally complex music was associated with enhanced integration within a “super-rich club” of hub regions. In contrast, ambient and traditional genres promoted more distributed and differentiated patterns of connectivity. Hub and centrality analyses further identified genre-specific effects on particular brain regions. While eight of the twelve genres elicited largely homogeneous network effects, two genres produced distinct regional signatures: Soft Jazz specifically engaged the left parieto-occipital region, whereas Progressive Instrumental Rock preferentially modulated the right posterior parieto-temporal region. These findings demonstrate that music listening evokes genre-dependent reorganization of information flow in functional brain networks, with only with only four genres recruiting a super-rich-club regime.

## 1 Introduction

Music is one of the most complex and time-consuming things that the human brain processes on a regular basis. It works in a highly coordinated way with auditory, motor, attentional, and affective systems(Koelsch [2014], Zatorre et al. [2007]). Listening to music has become a powerful naturalistic tool for studying how large-scale brain networks dynamically reorganize to meet ongoing processing demands, in addition to its perceptual and emotional aspects(Alluri et al. [2012], Toiviainen et al. [2020]). Continuous music, as opposed to controlled laboratory tasks, presents the brain with layered rhythmic, harmonic, and structural regularities that develop over seconds to minutes, necessitating ongoing integration across distributed cortical systems(Sachs et al. [2020]). A significant amount of research has delineated the regions that respond to music and the intensity of their co-activation. However, there is a paucity of knowledge regarding the routing of information through functional brain networks during listening, particularly whether different musical genres, which vary systematically in rhythmic complexity, predictability, and structural density, induce distinct regimes of directed communication. To fill this gap, we need to go beyond undirected correlational connectivity and use information-theoretic, directed network descriptions that can show the asymmetric, time-lagged dependencies that are common in auditory-cognitive processing.

Previous research in network neuroscience has established several foundational principles(Ahmadi et al. [2025], Saberi et al. [2025]. Large-scale brain networks frequently display non-random architectural features(Bullmore and Sporns [2012]), including small-world organization(Watts and Strogatz [1998]), modular structure(Meunier et al. [2010]), and hub-centric connectivity(Bullmore and Sporns [2009]), which facilitate efficient integration of distributed information. Transitioning from rest to task typically results in topological reconfiguration(Hearne et al. [2017], Krienen et al. [2014]), with hubs and core subnetworks exhibiting variable engagement based on processing demands(Cole et al. [2013]). Furthermore, directed measures of interaction, such as transfer entropy, are essential for elucidating how information is routed within the system (Schreiber [2000]), as these information-theoretic approaches can capture nonlinear, time-lagged dependencies and produce interpretable information-flow networks (Wibral et al. [2014], HaqiqiFar et al. [2025]). Collectively, these findings support the investigation of music listening as a genre-dependent alteration in the brain’s directed communication architecture, rather than solely as changes in spectral power or synchrony.

Despite these advances, a significant limitation persists: genre-specific information-flow networks remain challenging to compare and interpret reliably(Van Wijk et al. [2010]). In practice, transfer-entropy or other directed connectivity matrices are typically dense, sensitive to noise, and exhibit substantial inter-individual variability. As a result, downstream graph metrics are highly influenced by thresholding, denoising strategies, and other methodological factors(Ursino et al. [2020], Adamovich et al. [2022]). This dependence introduces a bottleneck for mechanistic interpretation, as differences in graph measures may reflect preprocessing choices rather than genuine neurophysiological reconfiguration. Additionally, although hub structure is frequently examined in the context of integrative cores, such as rich-club organization, the behavior of core topology in directed effective-connectivity networks during naturalistic listening remains unclear(Maggioni et al. [2021], Colizza et al. [2006]), as does the extent to which increased integration may entail trade-offs in network robustness(Albert et al. [2000]).

To address these challenges, this study analyzes a publicly available EEG dataset consisting of 128-channel recordings from 20 participants who listened, with eyes closed, to 12 musical excerpts representing distinct genres. Genre-specific directed information-flow networks are constructed by estimating transfer entropy between EEG channels at a fixed time lag, resulting in one directed adjacency matrix per participant and genre. To enhance interpretability and cross-subject reliability, a two-stage backbone extraction strategy is introduced, combining deep-learning-based denoising via a sparse autoencoder with an entropy-based stability filter that preferentially retains edges consistent across participants. Network density is selected using a topological sensitivity analysis to ensure stability. On these robust backbones, genre-dependent organization is quantified using compact topological fingerprints that summarize incoming and outgoing information flow patterns, as well as directed rich-club analysis normalized against degree-preserving null models. This approach is complemented by hub profiling, including measures of influence, brokerage, and virtual-lesion robustness, to link core topology with functional consequences.

This study contributes four key advances to network neuroscience in the context of naturalistic auditory cognition. Methodologically, a robust backbone-extraction pipeline for dense, directed transfer-entropy networks is introduced, combining sparse-autoencoder denoising with an entropy-based stability filter and density validation to reduce noise sensitivity and enable consistent cross-genre comparisons. Topologically, genre-specific information-flow fingerprints are derived to succinctly summarize how different musical structures redistribute incoming and outgoing information flow across the scalp-level network. Mechanistically, the analysis tests whether genres induce distinct directed rich-club regimes, linking changes in integrative core organization to differences in rhythmic or structural complexity and global coordination demands. At the network level, hub profiling, including PageRank and betweenness, is paired with virtual-lesion robustness analysis to determine whether increased integration is associated with greater fragility, thereby formalizing an efficiency–redundancy trade-off during demanding listening states. The following sections detail the dataset, transfer entropy estimation, and backbone construction used to obtain reliable genre-specific directed networks, report fingerprints and rich-club deviations that differentiate genres, and analyze hub roles and lesion-based resilience to interpret the functional consequences of these genre-tuned integration strategies for information-dynamic accounts of naturalistic music listening.

## 2 Results

### 2.1 Information Flow Networks and Topological Fingerprints

Directed information exchange between EEG channels during music listening is modelled by constructing genre-specific information flow networks using transfer entropy (TE). For each subject and genre, a full TE adjacency matrix was computed, where the edge weight from channel *i* to channel *j* quantifies the extent to which knowledge of the past signal of *i* improves prediction of the future signal of *j*, beyond what is predicted from *j*’s own history. TE was calculated as conditional mutual information with a time lag *u*. The estimator employed a *k*-nearest-neighbour (KSG-style) method, which estimates information-theoretic quantities by analysing the distances between neighbouring data points in the reconstructed state space. TE was evaluated at a time delay of *u* = 10 using a 5-nearest-neighbour configuration, yielding one directed TE matrix per subject and genre.

Due to the presence of weak or noisy edges and substantial inter-individual variability in dense TE matrices, a robust network backbone was extracted using a two-stage sparsification pipeline that combines deep learning denoising with an entropy-based stability filter. Initially, each raw TE matrix was processed through a sparse autoencoder that reconstructs the TE matrix while promoting a compact representation through an added sparsity constraint (reconstruction loss plus KL-based sparsity regularisation). This approach preserves dominant directed flow patterns and suppresses noisy, inconsistent connections. Subsequently, to retain only edges stable across subjects, the Shannon entropy of reconstructed edge weights *H*_*ij*_ was computed (lower entropy indicates more consistent edges across participants). A binary connectivity mask was then derived using a percentile-based threshold on *H*_*ij*_, and the final network density was fixed at 20%, resulting in a sparse graph that captures the most reliable backbone connections. The chosen sparsity level was justified by performing a density sensitivity analysis and tracking key topological indices (Fig.2A).

The network achieves full connectivity, with the giant component saturating at approximately 10% density and maintains small-world characteristics (*σ* ≈ 2.5 compared to random graphs) within a stable regime. This observation supports the selection of 20% density as a conservative threshold that preserves connectivity while avoiding excessive graph density (Fig.2A–C). At 20% density, the resulting adjacency matrix displays structured, non-random connectivity patterns (Fig. 1B), and the node-degree distribution exhibits a heavy tail on a log–log scale, consistent with scale-free-like organisation (Fig.2C).

**Figure 1:**
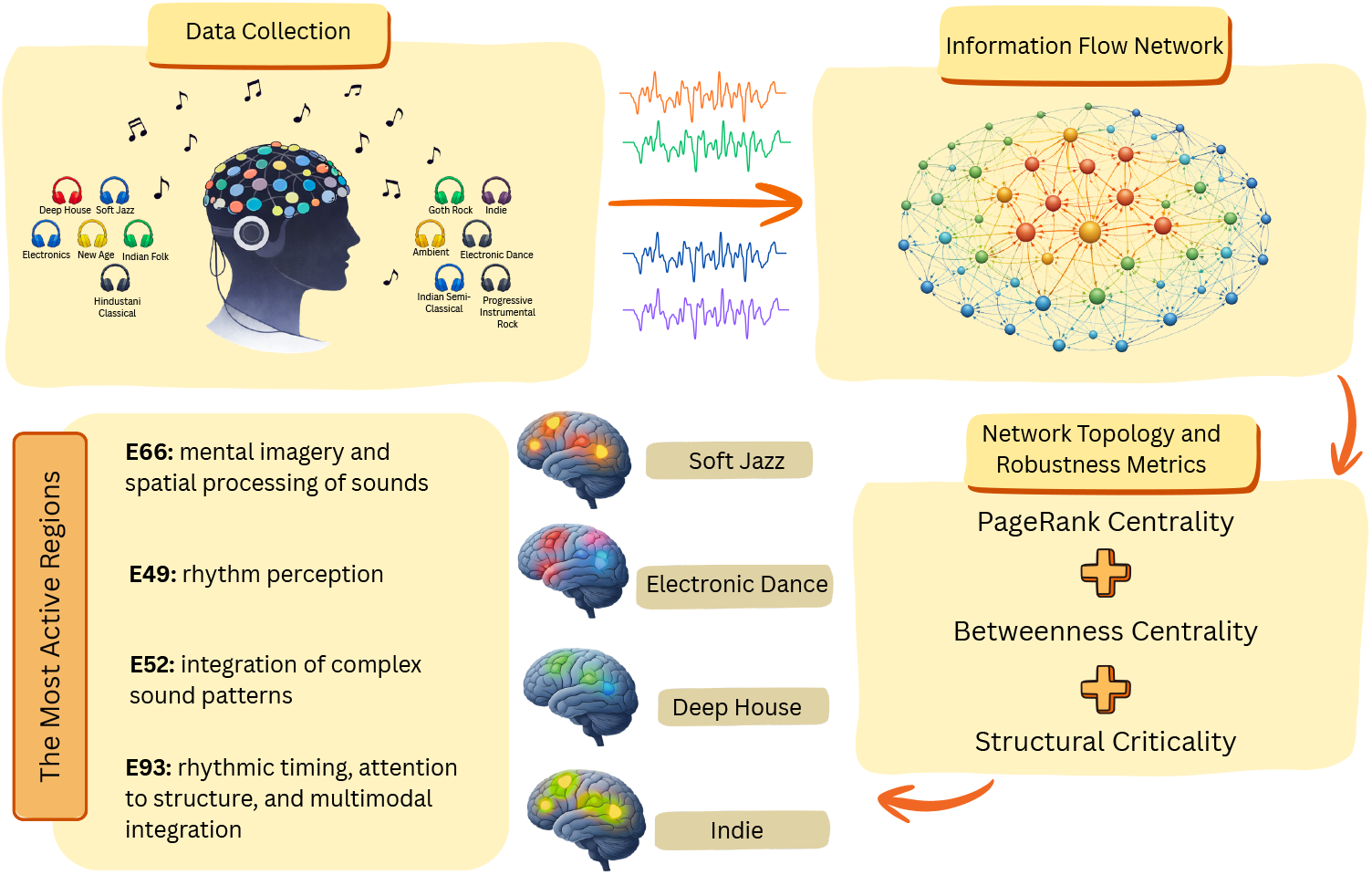
Schematic of the Proposed Methodology

**Figure 2:**
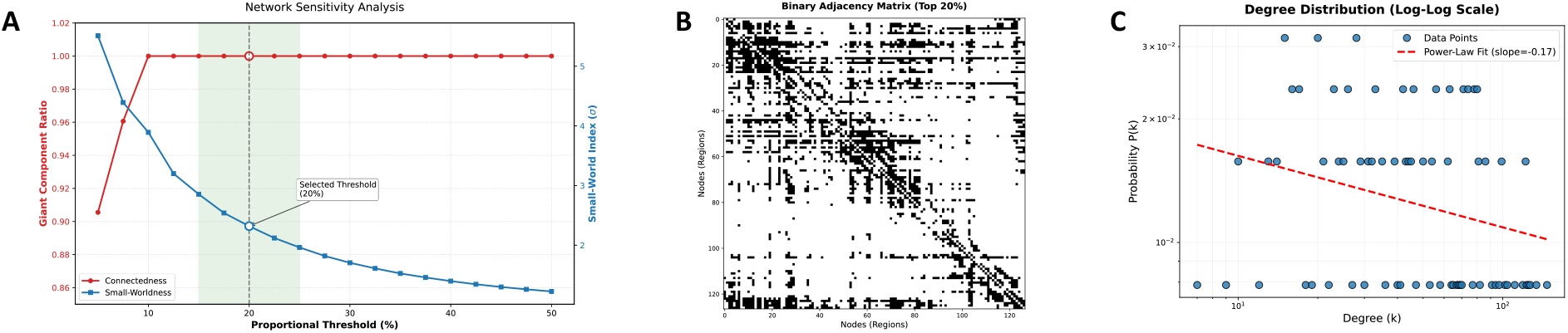
Topological validation and threshold selection. (A) sensitivity analysis and optimal threshold selection for the brain network. The plot illustrates the variations of the network’s topological indices over a density range of 5% to 50%. The red line represents the size of the Giant Component, indicating that the network reaches full connectivity at a density of 10%. The blue line shows the small-worldness index (*σ*), where the network, while ensuring full connectivity among all subjects, preserves small-world properties (*σ* ≈ 2.5) relative to random graphs. The green shaded region indicates the range of topological stability. (B) and (C) Structural network features were extracted at a 20% density. (B) Binary adjacency matrix: shows mean effective connectivity after thresholding. Black dots indicate significant connections, while white spaces show removed noise and weak connections. The row and column patterns indicate potential hub regions or major information pathways in the brain. (C) Node degree distribution: The probability distribution of node degrees, shown on a log-log scale, exhibits a linear downward trend and a heavy tail. These features indicate scale-free properties and confirm that the extracted structure reflects the intrinsic characteristics of biological networks.

Finally, to summarise genre-specific organisation in an interpretable way, we derived topological fingerprints from the reconstructed TE networks (Fig.3). For each genre, TE matrices were averaged across the 20 subjects (still evaluated at u=10u=10u=10, k=5), and node-wise entropy-based summaries of incoming vs outgoing information flow were visualised as scalp topographies (Fig.3A,D) alongside global average inflow entropy and average outflow entropy curves across genres (Fig.3B,C). These fingerprints provide a compact description of how each music genre reshapes the spatial distribution and the overall balance of information reception vs broadcasting within the brain’s directed functional network.

**Figure 3:**
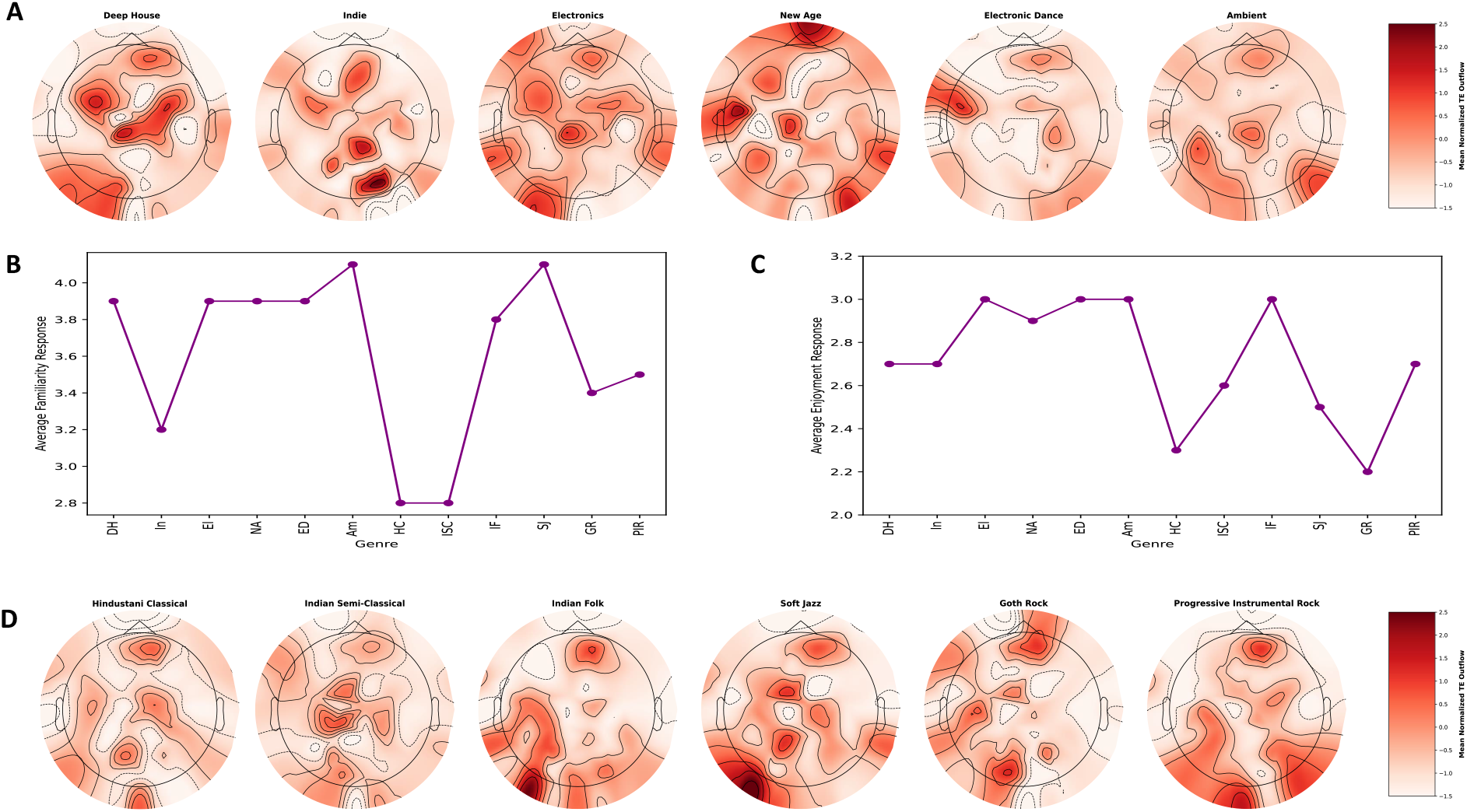
A, D) The average TE matrices were calculated for 20 subjects as they listened to 12 genres, measured across a time delay *u* = 10 and 5-nearest-neighbor. B) The plot shows the mean familiarity rating for each genre among participants, where 1 indicates the highest familiarity and 5 the lowest. C) The plot shows the mean enjoyment rating among participants, measured on a 1 to 5 scale, where 1 indicates the highest enjoyment and 5 the lowest.

### 2.2 Genre-Dependent Topological Reconfiguration

Based on the dynamic analysis of Rich-Club directed curves (*ϕ*_*norm*_), the brain network response to different music genres can be classified into three distinct topological patterns (Fig. 4). These patterns indicate how the brain adjusts the level of information integration based on the structural features of the music:

**Figure 4:**
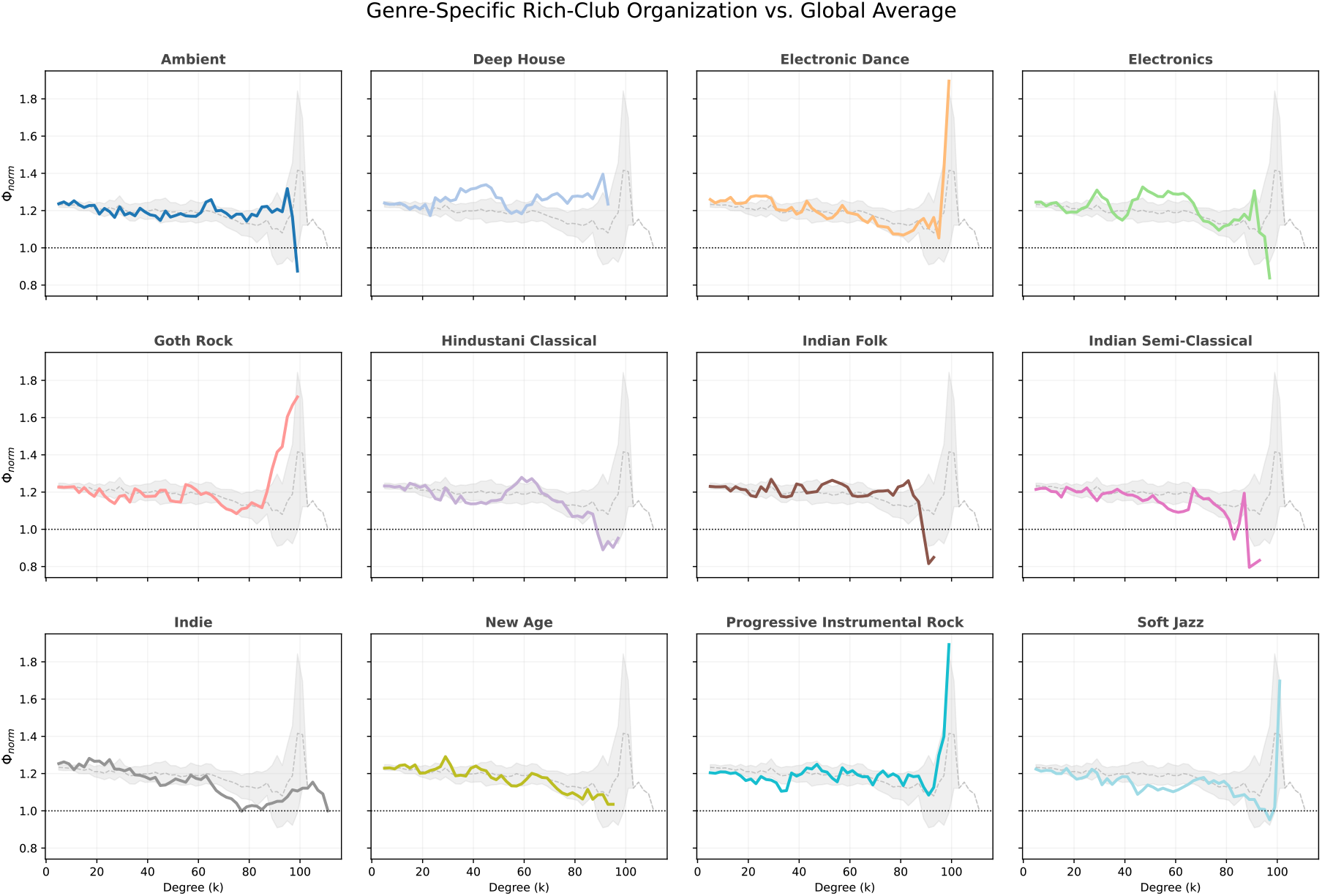
Genre-specific deviations of directed rich-club organization relative to the global profile. Small multiples show the normalized directed rich-club coefficient (Φ_*norm*_) for each genre (colored line) as a function of the total node degree (*k*). The gray-shaded region represents the mean ± standard deviation (SD) across all 12 genres, serving as a “reference structural profile.” The horizontal dashed line (*y* = 1) marks the random network boundary. Deviations of a genre’s curve from the gray region indicate changes in the brain’s information-integration strategy. Super-Integration: Genres such as Electronic Dance, Goth Rock, Progressive Instrumental Rock, and Soft Jazz show sharp, pronounced spikes at very high degrees (often *k >* 90). This rising pattern indicates the formation of highly dense, powerful connectivity cores among the network’s “super-hubs” (the most highly connected nodes) to support complex processing. Segregation: In contrast, genres such as Indian Folk, Indian Semi-Classical, and New Age show a decline at higher degrees, with the curves dropping below the global mean and even the random baseline. This indicates a reduced central role of hubs and a shift toward a more distributed, decentralized network topology for these music genres.

The Super Rich-Club Phenomenon: In Electronic Dance, Goth Rock, Progressive Instrumental Rock, and Soft Jazz genres, the Φ_*norm*_ curve undergoes a sudden and severe spike at very high degrees (*k >* 90) and reaches values much higher than the global average (and even higher than 1.6). This shift reflects a state of “hyper-integration,” where the “network elite” (the top 1–2% of hubs, likely including high-order auditory, motor, and executive regions) form an exceptionally cohesive and exclusive core. These genres typically have high rhythmic complexity, multiple sonic layers, or require precise temporal prediction. To process this volume of information and achieve precise synchronization, the brain mobilizes a “powerful central command,” despite the high metabolic cost of such dense, long-range connectivity. Stable Rich-Club Organization (The Standard Backbone): In the Deep House, Indie, New Age (partial), and Electronics genres, the curve fluctuates mostly above the dashed line (*y* = 1) and within the gray-shaded range (global average). They neither experience explosive spikes nor crashes. This pattern represents the preservation of the brain’s “Standard Backbone.” Here, the brain network strikes an optimal balance between integration and segregation. These types of music (like Deep House) usually have a regular, predictable structure that keeps the brain engaging enough to require central coordination of information but does not impose the excessive computational demand that necessitates the formation of a “Super-Core.”

Disrupted Core and Distributed Topology (Segregation): The curves for the Indian Folk, Indian Semi-Classical, Hindustani Classical, and Ambient genres drop sharply at high levels and fall below the random boundary line (*y* = 1). This means that the important brain nodes in this case are less connected than chance. This phenomenon represents a shift towards “distributed/segregated processing.” In this state, the central hubs downregulate their direct mutual coupling, and information processing becomes more localized within specific sensory modules. This shows that in music such as ambient (which is meant to be relaxing) or traditional music (which may have repetitive structures or cultural familiarity), the brain does not need to expend energy maintaining long-term, costly Rich-Club connections. This reduction in central integration may be associated with reduced focused awareness and states of mind-wandering. These results prove that the Rich-Club is not a fixed anatomical structure, but a dynamic functional arrangement. Like an intelligent manager, the brain only pays the high cost of forming a “super-core” (Super Rich-Club) when the complexity of the music demands it (e.g., Progressive Rock); otherwise (e.g., Ambient), it moves towards a more cost-efficient, distributed topology.

### 2.3 Genre-Tuned Rich-Club Regimes and Network-Level Consequences

The analysis of information dynamics within the brain network revealed that the brain’s topological structure during music listening is not a fixed anatomical feature but rather a fully dynamic functional substrate that undergoes fundamental changes in response to processing demands and the structural characteristics of the genre. Examination of the directed Rich-Club coefficient curves showed that the human brain can shift its information integration strategy across three distinct states.

First, in musical genres with high rhythmic and structural complexity, such as Electronic Dance, Progressive Instrumental Rock, and Soft Jazz, the brain network exhibited a phenomenon of super-integration, meaning the Rich-Club curves at high degree values showed abrupt increases, indicating the formation of highly dense and powerful communication cores among network hubs. Second, in genres with lower cognitive load and greater ambient structural features, such as Ambient and New Age, the network shifted toward a distributed topology, with reduced hub centrality and local information processing (Table.1).

**Table 1:**
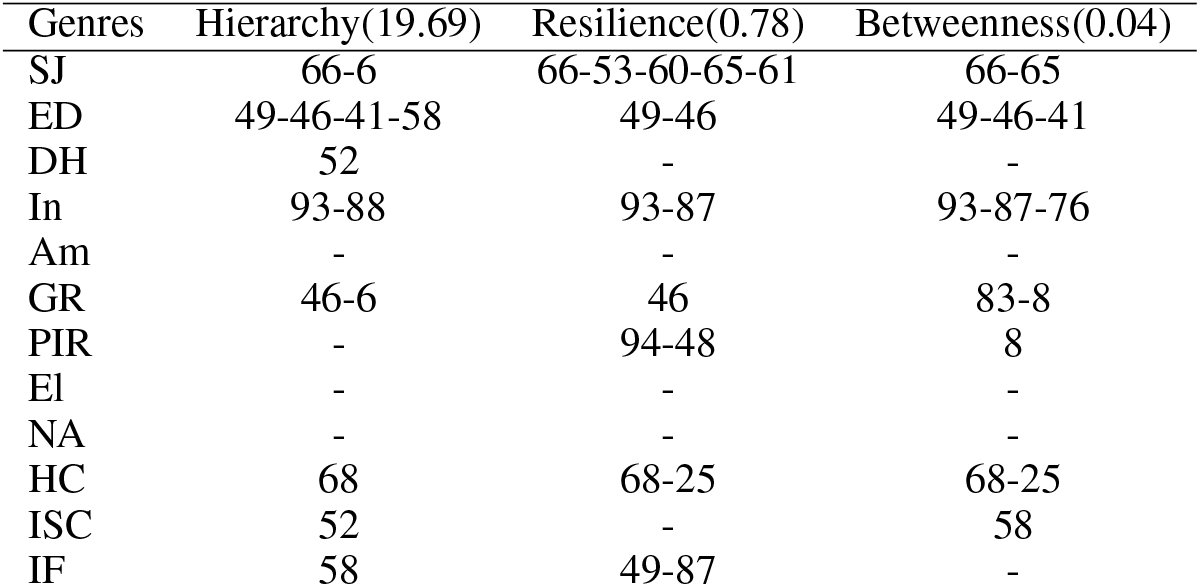
The Most Active Regions (after considering std)

To better understand the functional role of these structural cores, we analyzed the properties of network hubs in terms of hierarchical influence (PageRank), flow control (betweenness), and structural robustness (virtual lesion). PageRank centrality results showed that, across all musical genres, Rich-Club nodes consistently exhibited the highest network prestige, indicating that information flow preferentially converges on these nodes. In this sense, they play a central coordinating role in network dynamics. In contrast, betweenness centrality analysis revealed an additional layer of functionality. In complex genres such as Progressive Rock, network hubs act as communication bridges, linking distant brain modules and effectively shortening information processing pathways. These findings suggest that Rich-Club formation in such genres reflects the brain’s need for rapid integration of multisensory and temporally structured information.

Nevertheless, virtual lesion analysis revealed that this high level of efficiency comes at a substantial cost, namely, increased structural fragility. Comparison of resilience profiles across musical genres showed that the transition from calm music (Ambient) to high-energy music (Electronic Dance) corresponds to a shift from a resilient to a fragile network organization. In genres such as Electronic Dance and Soft Jazz, the virtual removal of a single top-ranked hub resulted in a pronounced drop in global network efficiency (exceeding two percent). This observation suggests that under cognitively demanding processing states, the brain concentrates its resources on a limited number of critical bottlenecks, effectively maximizing efficiency at the expense of redundancy. In contrast, in ambient genres, hub removal had only a minimal impact on overall efficiency, indicating a parallel distribution of process.

The spatial locations of the critical nodes identified in these analyses (e.g., E49, E94, and E66) reveal distinct processing patterns. In the Electronic Dance genre, which requires precise sensorimotor synchronization and beat prediction, the critical node E49 plays a central role. According to the electrode atlas, E49 is located in the left fronto-temporal region. This area is classically associated with the processing of spectral sound features, rhythm perception, and top-down auditory attention. The strong dependence of the network on this region suggests that, to maintain entrainment with fast and driving rhythms, the brain relies on concentrated activity within predictive circuits of the left temporal cortex.

In contrast, in the Progressive Instrumental Rock genre, which is characterized by unexpected changes in time signatures and complex harmonic structures, node E94 emerged as one of the most critical elements. This electrode is located in the right posterior parieto-temporal region. This area serves as a key hub for multisensory integration and higher-order auditory processing, particularly in the perception of complex changes in musical texture and auditory scene analysis. The activity of this hub as an informational bottleneck suggests that processing this genre requires the involvement of associative regions to link auditory perception with working memory and structural expectations.

Similarly, in the Soft Jazz genre, node E66 becomes structurally prominent. This electrode is located in the left parieto-occipital region. Although this area is typically associated with visual processing, within the context of music perception, particularly in improvisation-based music, activity in posterior parietal regions can be linked to mental imagery and spatial processing of sounds. The critical importance of this region in the Soft Jazz network may reflect the engagement of internal attention mechanisms and musical imagination that listeners use to follow fluid, non-repetitive melodic lines.

Finally, the striking differences in network topology between genres indicate that the “listening experience” is not a passive process. In Ambient and New Age genres, the activity of critical nodes is confined to anterior frontal regions (such as E25 and E124), but with much lower intensity and a more distributed influence. This pattern aligns with states of mind-wandering and reduced executive control, which are typically associated with feelings of relaxation and meditation. Overall, these findings support a model in which the brain acts as an intelligent resource manager, dynamically reconfiguring its connectivity topology in real time: forming an expensive yet efficient “super-core” to process rhythmic and structural complexity, and reverting to a distributed, low-cost organization in the absence of demanding processing challenges.

## 3 Discussion

The current study aimed to investigate whether specific musical genres induce significantly different patterns of directed communication in the human brain, and whether such reconfigurations can be consistently detected from scalp-level EEG after the reduction of dense transfer-entropy networks to a stable backbone across subjects. The findings yield a definitive and cohesive response. Directed information flow during naturalistic music listening is not consistent across genres nor accounted for by a singular integrative architecture; rather, the brain engages at least three distinct topological regimes. These range from a hyper-integrated super-rich-club core for rhythmically and structurally complex music (Electronic Dance, Goth Rock, Progressive Instrumental Rock, Soft Jazz), to a stable canonical backbone for moderately structured genres (Deep House, Indie, Electronics, New Age), and finally to a distributed configuration with sub-random hub coupling where hub coupling falls below chance for ambient and culturally familiar traditional music (Ambient, Hindustani Classical, Indian Semi-Classical, Indian Folk). This topological reorganization is accompanied by a systematic efficiency–fragility trade-off: the same hub concentration that enables rapid multisensory integration under high rhythmic load renders the network increasingly vulnerable to targeted disruption, with single-hub virtual lesions producing efficiency losses exceeding two percent in the most integrated genres. These observations reconceptualize naturalistic music listening not as a graded modulation of a static connectivity substrate, but as a genre-specific selection among qualitatively distinct information-routing strategies, each possessing unique computational advantages and structural expenses.

From a scientific perspective, the results support the interpretation of music listening as an active, demand-sensitive process in which the brain dynamically modulates directed integration. In this framework, communication is sometimes concentrated within a highly cohesive hub core for rhythmically and structurally complex genres, while in other cases, hub coupling is relaxed to promote distributed processing for more ambient or familiar structures. This perspective positions genre effects not solely as differences in oscillatory power, but as shifts in the routing of information through an integrative core.

From a methodological standpoint, this study recommends a pragmatic best practice for directed connectivity analyses. When networks are dense and noisy, as is often the case with transfer entropy (TE) matrices, interpretability is enhanced by selecting edges based on cross-subject stability and by justifying network density through topological sensitivity rather than convention. The integration of fingerprinting for interpretability with rich-club and lesion-based robustness for mechanism-level claims regarding integration and fragility offers a valuable template for application in other naturalistic paradigms.

In terms of translational implications, the observed efficiency–fragility trade-off suggests that directed network organization may serve as a quantitative metric for characterizing engagement or cognitive load during naturalistic listening. This has potential relevance for music-based interventions. However, due to the single-excerpt-per-genre design and limitations inherent to sensor-space analysis, these applications should be regarded as hypotheses rather than established biomarkers.

In conclusion, the brain does not process music through a static network. It listens with a reconfigurable one, always weighing the metabolic cost of making a dense, quickly integrating super-core against the strength of a distributed, segregated topology and choosing between them based on the rhythmic and structural needs of the stimulus. This study presents the first scalp-level, genre-resolved map of this arbitration by integrating a stability-aware backbone-extraction pipeline with directed rich-club analysis and virtual-lesion robustness profiling. It identifies a limited set of electrode-level bottlenecks—specifically, E49 for beat-driven entrainment, E94 for complex auditory-scene integration, and E66 for improvisation-linked mental imagery—whose functional significance appears to correlate with the computational demands of the genre. These results redefine music not only as an auditory stimulus but as a modifiable instrument for examining the brain’s directed communication framework, and they create a consistent basis for future source-localized, multi-excerpt, and clinical expansions to be effectively compared. The overarching implication is both conceptual and methodological: integration in the brain is not merely a characteristic; it is a decision—one that the brain continuously renegotiates, and one that music, due to its structured temporal complexity, is uniquely equipped to elucidate.

## 4 Materials and Methods

### 4.1 Data Acquisition and EEG Preprocessing

The dataset used in this study was provided by Miyapuram et al. [2021], which comprises EEG recordings with 128 channels collected from 20 Indian participants (16 men, four women) as they listened to 12 musical excerpts representing a wide variety of genres Pandey et al. [2021]. Note that each listening session involved a single song. The list of song genres used in the current research can be found in table 2.

**Table 2:** Song Genres Included in This Study.

| Genre List |  |  |  |  |  |
| --- | --- | --- | --- | --- | --- |
| Song ID | Genre | Short Form | Song ID | Genre | Short Form |
| 1 | Deep House | DH | 7 | Hindustani Classical | HC |
| 2 | Indie | In | 8 | Indian Semi-Classical | ISC |
| 3 | Electronics | El | 9 | Indian Folk | IF |
| 4 | New Age | NA | 10 | Soft Jazz | SJ |
| 5 | Electronic Dance | ED | 11 | Goth Rock | GR |
| 6 | Ambient | Am | 12 | Progressive Instrumental Rock | PIR |

At the start of a trial, participants were asked to close their eyes after hearing a short beep, and the song was then played through speakers. When a double beep signaled the end of the song, they opened their eyes and rated how familiar they were with the piece and how much they enjoyed it. Both ratings were given on a five-point scale, where one indicated the highest familiarity or enjoyment, and five the lowest. However, emotional engagement was not investigated in our article, as it was beyond the scope of this study.

The raw EEG recordings (referred to as sourcedata) include complete sessions for each participant, beginning with a two-minute resting baseline, followed by neural responses to all 12 songs, which were presented in a randomized order. Each song was presented once per participant. The raw and segmented EEG datasets have been organized according to the BIDS standard using the eeg2bids plugin in EEGLAB. For preprocessing, the EEG signals were processed in EEGLAB Delorme and Makeig [2004] following the procedure described by Pandey et al. [2021]. A high-pass linear FIR filter at 0.2 Hz was applied, and CleanLine Mullen [2012] was used to remove 50 Hz power-line noise. The recordings were downsampled to 250 Hz and divided into 12 trials corresponding to each song. Noisy channels were automatically detected using spectrum-based criteria (3 SD threshold) and removed. Artifact removal was conducted using the Multiple Artifact Rejection Algorithm (MARA) Winkler et al. [2011], a machine learning-based independent component analysis method. Removed channels were interpolated using spherical interpolation, and the data were re-referenced to the common average.

### 4.2 Transfer Entropy and Measuring Information Flow

Shannon entropy provides a fundamental measure of uncertainty in information theory. Consider a random variable *X*, with outcome *x*, and a probability distribution *p*(*x*). The central idea of Shannon information Shannon [1948] is that observing a particular outcome *x* reduces the uncertainty about the system.

Shannon proposed using a monotonic transformation of the probability to quantify this reduction in uncertainty. The information gained from observing an outcome *x* with probability *p*(*x*) is therefore defined as:

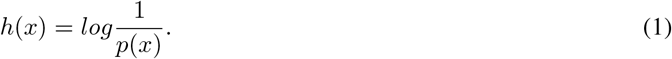

This definition makes it immediately clear that unlikely outcomes convey greater information when they occur. Shannon entropy *H*_*x*_ is defined as the expected information content associated with repeated realizations of *X*. In other words, it represents the average uncertainty reduced upon observing the outcomes of *X*.

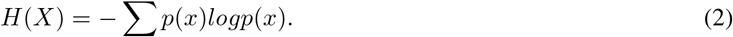

For continuous values, on the other hand, Shannon entropy can be defined as:

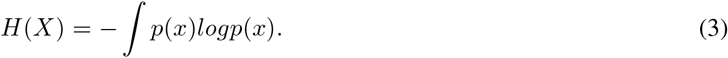

Shannon entropy helps us to understand how unpredictable or “informative” a system is. It reminds us that the more uncertain something is, the more information we gain when we finally know its outcome.

Transfer Entropy, on the other hand, serves as a powerful quantitative measure for capturing the directional flow of information between dynamic systems, representing a model-independent realization of Wiener’s formulation of observational causality. Wiener’s framework asserts that, given two concurrently measured processes *X* and *Y*, process *X* can be regarded as causal with respect to *Y* if incorporating the past states of *X* yields a superior prediction of *Y* compared to predictions derived solely from the past states of *Y*. Another aspect of Wiener’s principle posits that the past state of process *X* is considered informative about the future state of *Y* if it provides additional predictive value beyond what is already contained in the past state of *Y*. The latter principle can be written as follows Wibral et al. [2014]:

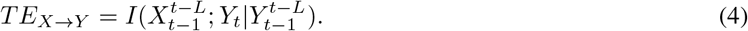

Note that *T E*_*X*→*Y*_ represents the directional flow of information from process *X* to process *Y*. Moreover, 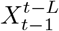 is a vector of past values of *X*, starting from time *t* − 1 going back L stepts: *X*_*t*−1_, *X*_*t*−2_, …, *X*_*t*−*L*_, an analogous statement applies to 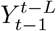. *Y*_*t*_, on the other hand, represents the future value of *Y* that we aim to predict using the past of both *X* and *Y*. *I*(.;.|.) is the conditional mutual information. In other words, it measures how much information *X*’s past provides about *Y*_*t*_ given that we already know *Y* ‘s past.

### 4.3 KSG Technique

Mutual information quantifies the amount of shared information between two random variables, capturing both linear and nonlinear dependencies. Unlike correlation, mutual information detects any form of statistical dependence, making it a powerful tool for understanding complex relationships in data. Mutual information measures how much knowing one random variable reduces the uncertainty about the other Kozachenko [1987], and can be estimated by:

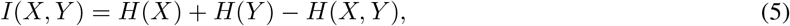

which implies that the errors arising from the individual estimates would, in general, not cancel out. For this reason, a different approach is adopted Kraskov et al. [2004]. In what follows, two closely related algorithms are introduced, both grounded in the aforementioned idea. In both cases, we employ the maximum norm for two points *z*_*i*_ = (*x*_*i*_, *y*_*i*_) and *z*_*j*_ = (*x*_*j*_, *y*_*j*_) in the space *Z* = *X* × *Y*, the maximum norm is defined as:

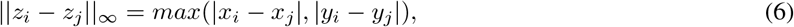

while any norms can be used to measure ||*x*_*i*_ − *x*_*j*_|| and ||*y*_*i*_ − *y*_*j*_ ||, these norms need not to be identical. For convenience, we adopt the following notation from previous works Kraskov et al. [2004]; let *ϵ*_*i*_*/*2 denote the distance from *z*_*i*_ to its *k*-th nearest neighbor, and let *ϵ*_*x,i*_*/*2 and *ϵ*_*y,i*_*/*2 denote the corresponding distances between the same points projected onto *X* and *Y* subspaces, respectively. It then follows that:

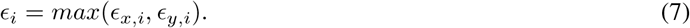

As demonstrated by Kraskov et al. [2004], the Shannon entropy is defined as:

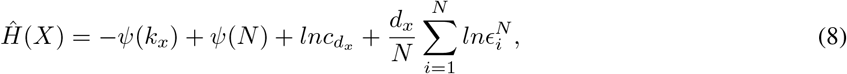

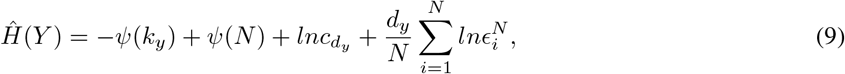

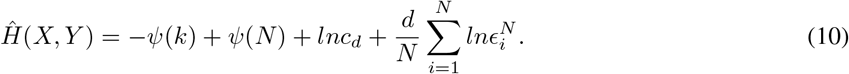

The above-mentioned formulas can be derived from the Shannon entropy equation, and the probability distribution *p*_*k*(*ϵ*)_. Also note that: 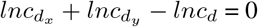, where *d*_*x*_ and *d*_*y*_ denote the dimensionality of *x* and *y*, respectively, and *c*_*d*_ represents the volume of the unit ball in *d*-dimensional space. This is also accurate for 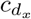 and 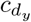. Furthermore, *ψ*(*x*) is digamma function which represents 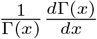. Also, *N* refers to the number of data samples used to estimate mutual information. Therefore, mutual information can be expressed as follows:

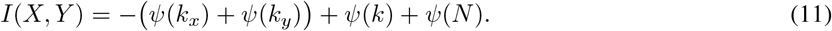

The former equation is valid for any choice of *k*, which means it is unnecessary to fix a particular *k* when estimating the marginal entropies. If there are in total *n*_*x*_(*i*) points within the vertical strip defined by 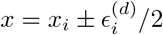, then 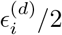 corresponds to the distance to the (*n*_*x*_(*i*) + 1)-th neighbor of *x*_*i*_. Therefore, the entropy can be written as:

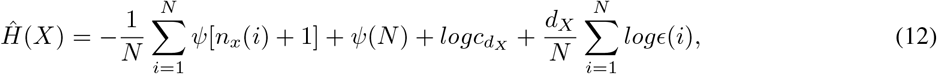

hence, mutual information can be expressed as:

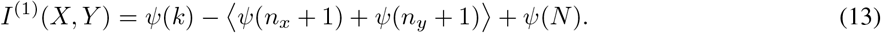

Alternatively, instead of counting neighbors *n*_*x,i*_ and *n*_*y,i*_ by fixed k-nearest distances, the number of points lying within distances *ϵ*_*x,i*_*/*2 and *ϵ*_*y,i*_*/*2 from point *i* is counted. The MI estimator is then computed using these counts, reflecting the local neighborhood sizes adapted to the marginal spaces, improving bias and variance properties.

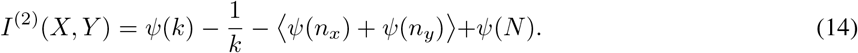

The Kraskov–Stögbauer–Grassberger (KSG) estimator provides a robust, bias-minimized method for computing mutual information using k-nearest neighbor statistics. Its adaptive nature makes it highly efficient and accurate, even for small sample sizes and complex, non-Gaussian data distributions.

### 4.4 Transfer Entropy Estimator

Transfer entropy can be regarded as conditional mutual information over time, and conditional mutual information can be directly calculated by Shannon entropy. From Eq.1 and Eq.2, it can be concluded that:

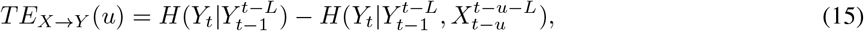

where *u* represents the time lag between the source process *X* and the target process *Y*. With the Kraskov-Grassberger-Stögbauer approach, the expression becomes:

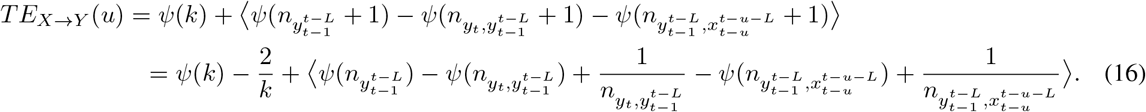

The transfer entropy estimator shows how one time series can influence another over time by building on the idea of mutual information. By looking at the past states of both the source and target, it gives a practical way to measure the flow of information and uncover causal relationships in complex systems.

### 4.5 Network Sparsification and Backbone Extraction

To separate stable functional connections from spurious noise and inter-individual variability, we used a sparsification framework based on deep learning, combined with an information-theory-based filter. The raw transfer entropy matrices were first processed with a sparse autoencoder. A sparse autoencoder is an autoencoder that is encouraged to use only a small subset of connections, highlighting the most important connections rather than retaining all of them. The autoencoder map is defined as:

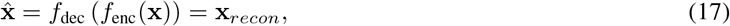

where *f*_enc_ is the encoder, *f*_dec_ is the decoder, and 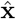 is the reconstructed TE matrix.Unlike other standard dimension-reduction techniques, the sparse autoencoder we used was trained using a loss function that included mean-squared error (MSE) for reconstruction and an L1 regularization penalty on the hidden-layer activations:

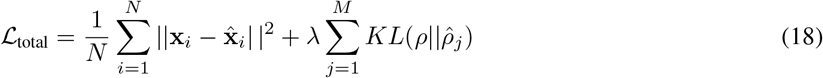

The hyperparameter *λ* is used to control the sparsity level of the network and *KL* is the Kullback-Leibler divergence that constrains the average activation of regions 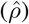 to remain close to a low target value (e.g., *ρ* = 0.05).The sparsity constraint forced the model to learn a compact latent representation of brain networks and effectively denoise the data by prioritizing the most salient connectivity patterns shared across participants. As a result, reconstructed matrices preserved the main directional information flow, while exhibiting reduced noise. Using the entropy-based filter and generating masks after the reconstruction step, the binary connectivity masks for each music genre were produced. This process involved evaluating the stability of each edge across subjects. To this end, the Shannon entropy of the reconstructed weights was calculated to minimize their variability. Then Shannon entropy is calculated by:

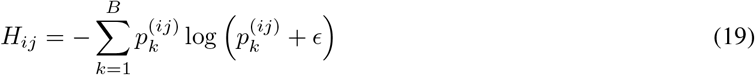

where *H*_*ij*_ is the entropy score for edge (*i, j*) and *ϵ* is a small constant. The sparsity constraint forced the model to learn a compact latent representation of brain networks and effectively denoise the data by prioritizing the most salient connectivity patterns shared across participants. As a result, reconstructed matrices preserved the main directional information flow, while exhibiting reduced noise. Using the entropy-based filter and generating masks after the reconstruction step, the binary connectivity masks for each music genre were produced. This process involved evaluating the stability of each edge across subjects. To this end, the Shannon entropy of the reconstructed weights was calculated to minimize their variability. A binary mask is built for each genre using the Percentile-Based Threshold for edge selection as:

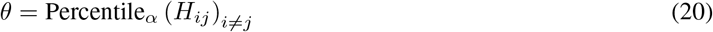

where *θ* is the threshold for the *α*_*th*_ percentile of all *H*_*ij*_. This dual-filtering approach resulted in sparse binary masks that preserved the network backbone; that is, a subset of connections that were both structurally meaningful and consistent across participants. This ensures that our findings reflect robust neural mechanisms rather than transient noise.

### 4.6 Genre-Dependent Modulation of Core Topology

The “Rich-Club” phenomenon, introduced by Van Den Heuvel and Sporns [2011], refers to the tendency of high-degree nodes (network hubs) to form dense, non-random connections, leading to a coherent central infrastructure for integrating information within the network. To investigate the tendency of drivers to form dense connections with each other, we generalized the Rich-Club coefficient to directed graphs. Unlike conventional analyses of undirected graphs, this study preserved the direction of information flow. Let *G* = (*V, E*) be a directed graph with *N* nodes and *E* edges. For each node *i*, the total degree (*k*_*tot*_) is defined as the sum of the input and output degrees:

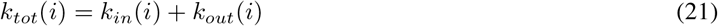

Let *N*_*>k*_ represent the number of nodes in this club. The directed rich club coefficient, Φ(*k*), is the ratio of directed edges among club members (*E*_*>k*_) to the maximum possible edges between them. In a self-loop directed graph, this maximum is *N*_*>k*_(*N*_*>k*_ − 1). Therefore:

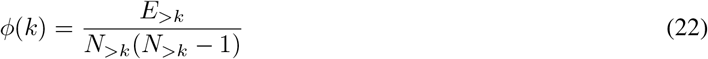

where *E*_*>k*_ is the number of directed edges (*u, v*) such that both *u* and *v* belong to *V*_*>k*_.

Since high-degree nodes naturally have a higher chance of being randomly connected, the coefficient Φ(*k*) needs to be normalized to a random model (Null Model). To this end, we generated a set of 100 directed random networks (*G*_*rand*_) for each observed network. These networks were generated using an edge-rewiring algorithm such that the distributions of input degree (*k*_*in*_) and output degree (*k*_*out*_) for all nodes were exactly the same as in the original network, but the global topological structure was destroyed. The normalized Rich-Club coefficient (Φ_*norm*_) was calculated as follows:

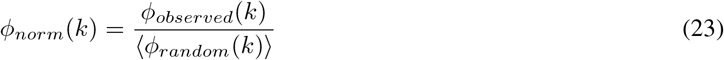

where ⟨Φ_*random*_(*k*)⟩ is the average of the coefficients calculated for the set of random networks. The value Φ_*norm*_(*k*) *>* 1 indicates the existence of a Rich-Club organization beyond the random limit; that is, the network hubs have a significant tendency to communicate with each other and form a dense core.

